# Morph bias in inflorescences and individual plants reduces opportunities for geitonogamy in a monomorphic enantiostylous species

**DOI:** 10.64898/2026.01.29.702576

**Authors:** S. B. Rhuthuparna, Vinita Gowda

## Abstract

- Evolution of enantiostyly, a stylar polymorphism characterised by left– or right-handed flowers, is predicted to counter selfing through disassortative pollen movement. However, in monomorphic enantiostyly, both morphs are present on the same individual, and geitonogamy has been proposed to depend on the morph ratio of an individual.
- Using the monomorphic enantiostylous *Didymocarpus podocarpus* (Gesneriaceae), endemic to the Eastern Himalayas, we hypothesised that geitonogamous events would be reduced when plants have a biased morph ratio, due to disassortative pollen movement. We first established the natural morph ratios and reproductive compatibilities of the morphs, and next examined the role of morph ratios in pollen movement by recording pollinator visitation patterns using visual observations and quantum dots.
- Our results show that most inflorescences and plants exhibit a stochastic skew in their morph ratios, while populations maintain an isoplethic ratio. Pollen transfer and pollination success were higher between morphs than within morphs, and inter-morph pollinator switches within a plant significantly decreased with an increase in morph bias.
- We show that within a plant, biased morph ratios combined with disassortative pollen movement can limit the occurrence of geitonogamous events, thus providing a mechanism to improve pollination success while also retaining reproductive assurance.

## Introduction

In hermaphroditic flowers, the spatial arrangement of male and female reproductive structures can affect their reproductive success (Webb & Lloyd, 1986; Richards, 1997; Barrett *et al*., 2000; Barrett, 2002; Li *et al*., 2009; Barrett, 2010). Hermaphroditic plants with inter-mating floral morphs, such as plants with stylar polymorphisms, are predicted to have evolved to counter selfing and to promote outcrossing (Barrett et al., 2000; Barrett & Harder, 2005; Pauw, 2005; Barrett, 2010; De Almeida & de Castro, 2019). In species with stylar polymorphisms, on a population scale, pollination and its outcome depend on the number of flowers of each floral morph that are displayed simultaneously within and between plants (Waites & Ågren, 2004; Brys & Jacquemyn, 2010). For instance, in stylar polymorphism like heterostyly, individual plants display only one type of floral morph. This facilitates pollen movement among individuals carrying dissimilar morphs, that is, inter-morph cross-pollination, thus ensuring higher outcrossing rates within a population (Keller *et al*., 2014). However, in enantiostyly, which represents a distinct type of stylar polymorphism characterised by left (L-morph) or right-handed (R-morph) flowers, the morphs can be present on the same individual or segregated between individuals (Table 1). Enantiostylous flowers are identified by left or right-handed orientation of the style (non-reciprocal enantiostyly) or both style and stamen (reciprocal enantiostyly) away from the main floral axis. The two morphs may be present in two separate individuals, resulting in dimorphic enantiostyly (DE), or within the same individual, as observed in monomorphic enantiostyly (ME). A majority of the ME species are reported to be self-compatible (Table 1; Wang et al., 1995; Gao et al., 2006; Tang & Huang, 2005; Ren et al., 2013; Richman & Venable, 2018; Mora-Carrera et al., 2019; Paudel *et al.,* 2024; Johnson *et al*., 2025) and thus, the presence of both the morphs within an individual can pose a risk of geitonogamous pollen transfer.

**Table 1.**
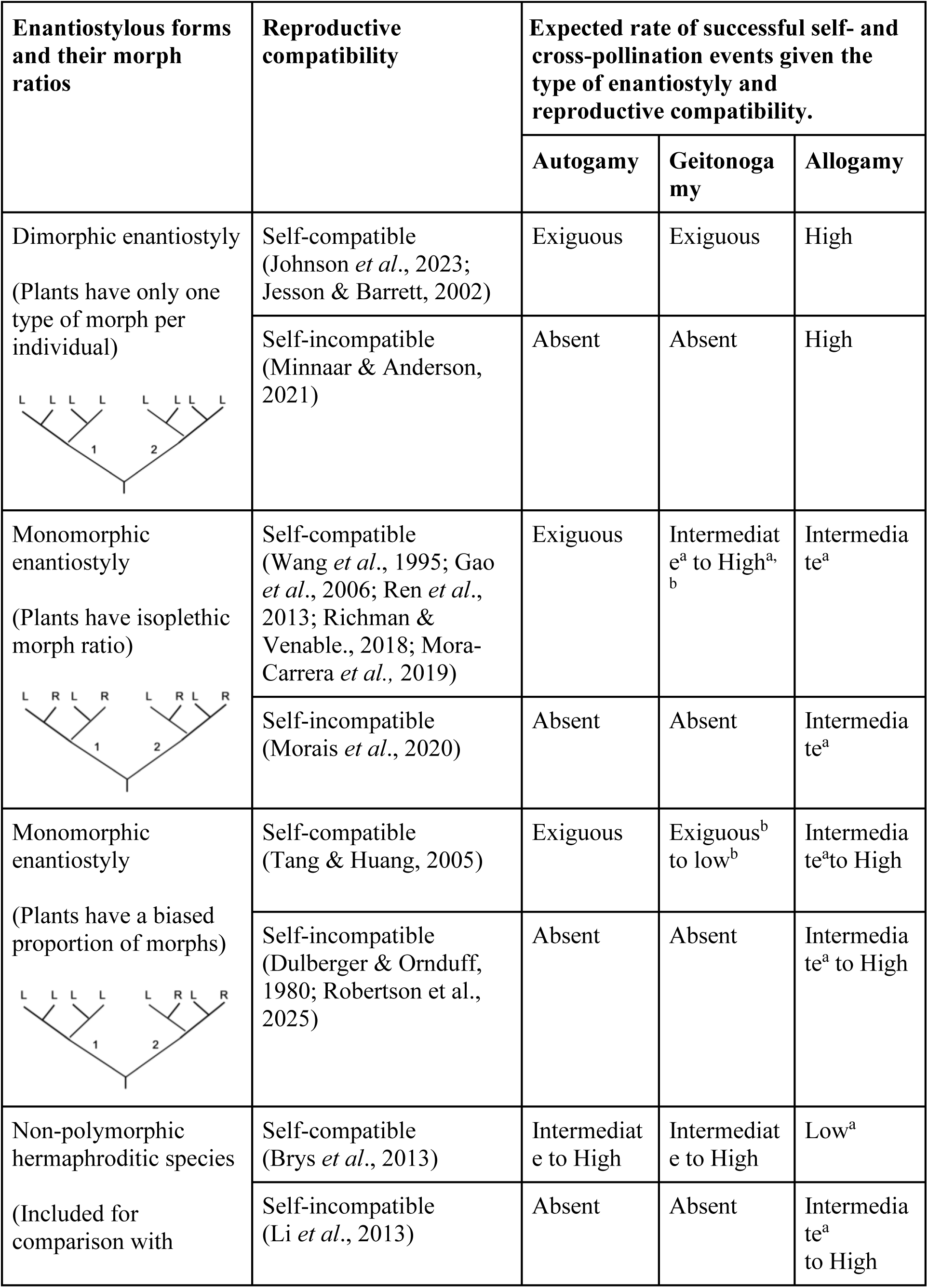
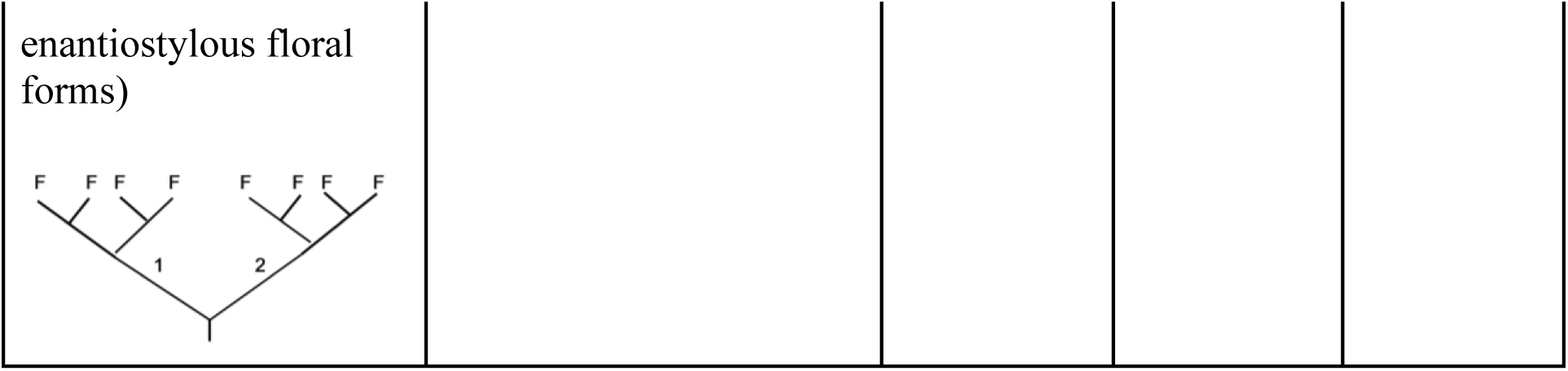
The potential for successful self-pollination (autogamy and geitonogamy) and cross-pollination (allogamy) events in different types of enantiostylous plants. Floral arrangement of L morph (L), R morph (R), and hermaphroditic flowers (F) within two inflorescences (1 and 2) and within a plant are illustrated. The pollination evaluation is based on the morph ratios characteristically known for the specific type of enantiostyly and their known reproductive compatibility (references included). The superscripts refer to pollination outcomes under two specific conditions: a–when within-plant pollen loss is expected, b–when floral arrangement can affect pollen movement within an inflorescence on a plant.

| Enantiostylous forms and their morph ratios | Reproductive compatibility | Expected rate of successful self- and cross-pollination events given the type of enantiostyly and reproductive compatibility. |  |  |
| --- | --- | --- | --- | --- |
|  |  | Autogamy | Geitonogamy | Allogamy |
| Dimorphic enantiostyly<br>(Plants have only one type of morph per individual) | Self-compatible (Johnson *et al.*, 2023; Jesson & Barrett, 2002) | Exiguous | Exiguous | High |
|  | Self-incompatible (Minnaar & Anderson, 2021) | Absent | Absent | High |
| Monomorphic enantiostyly<br>(Plants have isoplethic morph ratio) | Self-compatible (Wang *et al.*, 1995; Gao *et al.*, 2006; Ren *et al.*, 2013; Richman & Venable., 2018; Mora-Carrera *et al.*, 2019) | Exiguous | Intermediate<sup>a</sup> to High<sup>a, b</sup> | Intermediate<sup>a</sup> |
|  | Self-incompatible (Morais *et al.*, 2020) | Absent | Absent | Intermediate<sup>a</sup> |
| Monomorphic enantiostyly<br>(Plants have a biased proportion of morphs) | Self-compatible (Tang & Huang, 2005) | Exiguous | Exiguous<sup>b</sup> to low<sup>b</sup> | Intermediate<sup>a</sup> to High |
|  | Self-incompatible (Dulberger & Ornduff, 1980; Robertson *et al.*, 2025) | Absent | Absent | Intermediate<sup>a</sup> to High |
| Non-polymorphic hermaphroditic species<br>(Included for comparison with | Self-compatible (Brys *et al.*, 2013) | Intermediate to High | Intermediate to High | Low<sup>a</sup> |
|  | Self-incompatible (Li *et al.*, 2013) | Absent | Absent | Intermediate<sup>a</sup> to High |

Floral handedness in enantiostylous species has been shown to enhance the movement of pollen between flowers of different morphs, resulting in a disassortative pollen movement (i.e., L to R or R to L), thus reducing self-pollen transfer (geitonogamy) when compared with plants that lack enantiostyly (Table 1; Richards, 1997; Barrett *et al*., 2000; Jesson *et al*., 2003; Jesson & Barrett, 2005). Between DE and ME types of enantiostyly, geitonogamy is lowest in DE species, whereas ME species may exhibit intermediate levels of geitonogamy (Jesson & Barrett, 2005; Saltini *et al*., 2025). Thus, rates of geitonogamy may vary with the type of enantiostyly, where the main differentiating factor is the distribution of L and R morphs within an individual.

Among ME species, a population may consist of individual plants with an equal distribution of L and R flowers (Wang *et al*., 1995; Gao *et al*., 2006; Martins, 2008; Ren *et al*., 2013; Richman & Venable, 2018; Mora-Carrera *et al*., 2019; Braga *et al*., 2022) or a skewed proportion of morphs, where individual plants exhibit partial or complete bias towards either one of the morphs (Dulberger & Ornduff, 1980; Tang & Huang, 2005; De Almeida *et al*., 2018; Robertson *et al*., 2025). Since ME species bear both L and R morphs on the same individual plant, both the visitation sequence of pollinators within an individual as well as the relative abundance of L and R morphs are predicted to contribute towards geitonogamous pollen transfer (Table 1; Jesson & Barrett, 2005; Tang & Huang, 2005; Mora-Carrera *et al*., 2019; Barrett & Fairnie, 2024). However, in ME individuals with an equal number of L and R morphs, that is, an isoplethic morph ratio, geitonogamous pollen transfer may be inevitable since both active morph-switches and random switches by pollinators will result in high geitonogamous pollen transfer. Thus, a reduced opportunity for geitonogamous pollen transfer may be achieved in an ME species if morph ratios deviate from an isoplethic ratio within an individual. A highly biased morph ratio where one morph is dominant over the other within a plant can ensure that even under random pollinator movement within an individual, the disassortative pollen movement (L to R and R to L) is mostly between individuals.

To the best of our knowledge, the effect of the relative abundance of morphs within the plant on opportunities for geitonogamy in natural populations of ME plants remains to be tested. Our present research aims to fill this void by investigating the ME species *Didymocarpus podocarpus* (Gesneriaceae), an understorey, perennial, mostly lithophytic herb restricted to the eastern Himalayas. We carried out *in situ* studies on two populations of *D. podocarpus* and tested the hypothesis that geitonogamous events will be reduced when plants have a biased morph ratio, especially due to disassortative pollen movement between morphs. Though ME has been reported from multiple genera within Gesneriaceae (Harrison *et al*., 1999; Gao *et al*., 2006; Ling *et al*., 2020; Prasanna, 2023), enantiostyly remains an understudied topic. The study was conducted across two populations of *D. podocarpus* to ensure that the morph ratio is not a population-specific character.

The aims of our investigation were as follows: (1) to quantify monomorphic enantiostyly by documenting morph ratios in natural populations across multiple years; (2) to perform hand-pollination experiments to establish autonomous self-pollination, and to check intra– and inter-morph compatibility; (3) to evaluate pollen transfer patterns by the pollinators in the two morphs, and (4) to finally integrate morph ratios, reproductive compatibilities of the two morphs, and pollinator visitation patterns to infer the potential of geitonogamy within a plant. Based on these objectives, we address the following key questions: (i) Do the morph ratios within a plant and a population deviate from an isoplethic ratio? (ii) Do self and cross treatments yield similar reproductive success (fruit set and seed set) in the two morphs? (iii) Do morph ratio and pollinator visitation patterns within an individual affect opportunities for geitonogamous pollen transfer?

## Materials and methods

### Study species and study sites

*Didymocarpus podocarpus* (Gesneriaceae) is a perennial herb distributed in the Himalayan foothills of Northeast India, Nepal, Bhutan, and China. It is mostly lithophytic (occasionally terrestrial), found on moss-covered rocks. The species flowers between late July and early September, and the flowering within an individual plant lasts for a maximum of 15–20 days. The plants produce 1–9 cymose inflorescences, each bearing ≤ 11 open flowers at the time of census (mean ± SE number of open flowers per inflorescence is 2.84 ± 0.11 and per individual plant is 6.28 ± 0.41). The species exhibits monomorphic, reciprocal enantiostyly with distinct L and R morphs (Fig. 1; Fig. S1). The flowers are tubular, have a nectary at the base of the tube and last for 5 days. Stigma receptivity did not differ between the two morphs across their floral longevity, based on the hydrogen peroxide test (Kearns & Inouye, 1993). All experiments were conducted in two natural populations of *D. podocarpus* within India, the Takdah Reserve Forest, West Bengal (27°01’58” N, 88°20’23” E, elevation 1812m), and Gangtok, Sikkim (27°20’30” N, 88°37’16” E, elevation 2001m), during August in the years 2022, 2024, and 2025. We studied a total of 153 plants for the estimation of morph ratio (section 2.2), out of which 72 were used in quantifying pollinator visitation sequence within individual plants (section 2.6). Out of 153, only 109 plants were available to measure fruit set. Since experimentation on this species in shade house conditions is challenging due to both collection restrictions from the wild and survivability rates in the shade house, all studies were carried out in the wild populations. The primary pollinator (98.8% visits) of *D. podocarpus* was identified as *Bombus breviceps* (Apidae; Williams, 2022), and throughout this study, the term ‘*Bombus* bee’ refers only to this species.

**Fig. 1.**
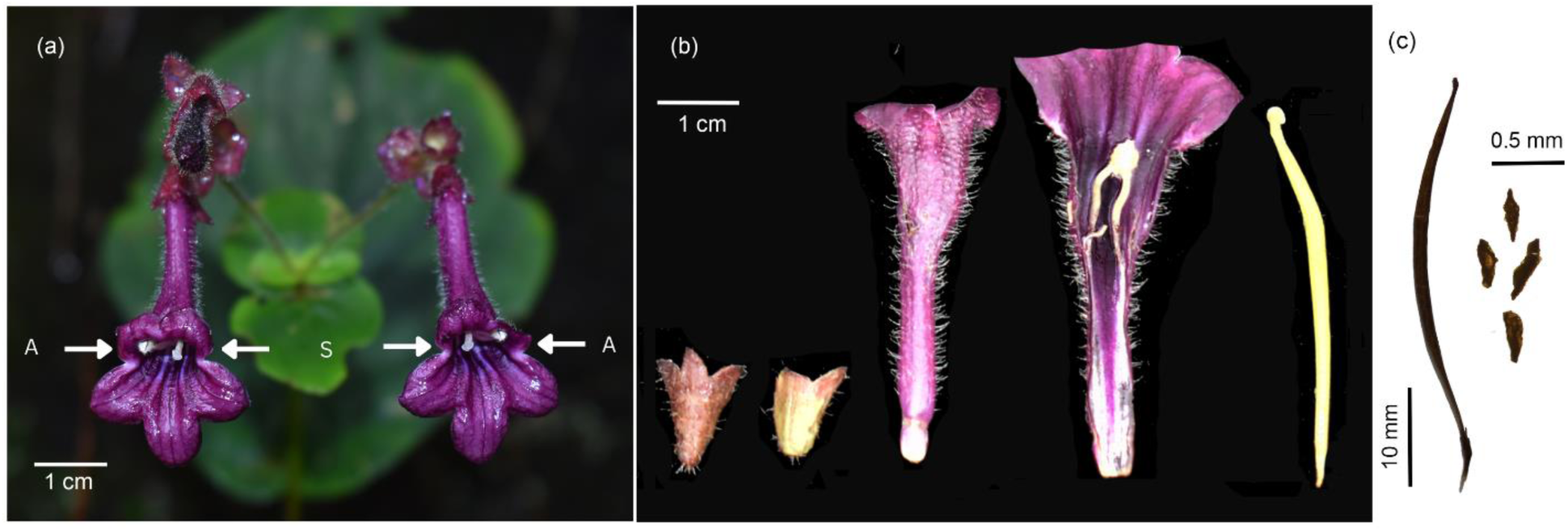
Floral and fruit morphology of enantiostylous *D. podocarpus*. (**a**) From left to right of the image – L morph and R morph. Arrows point to the reciprocal placement of anther (A) and stigma (S) on the L and R morphs. (**b**) Floral dissection (from left to right) – adaxial surface of calyx, abaxial surface of calyx, abaxial surface of corolla tube, abaxial surface of corolla tube showing fusion of the anther to the corolla tube, and pistil. (**c**) mature fruit, and a microscopic image of seeds.

**Fig. 2.**
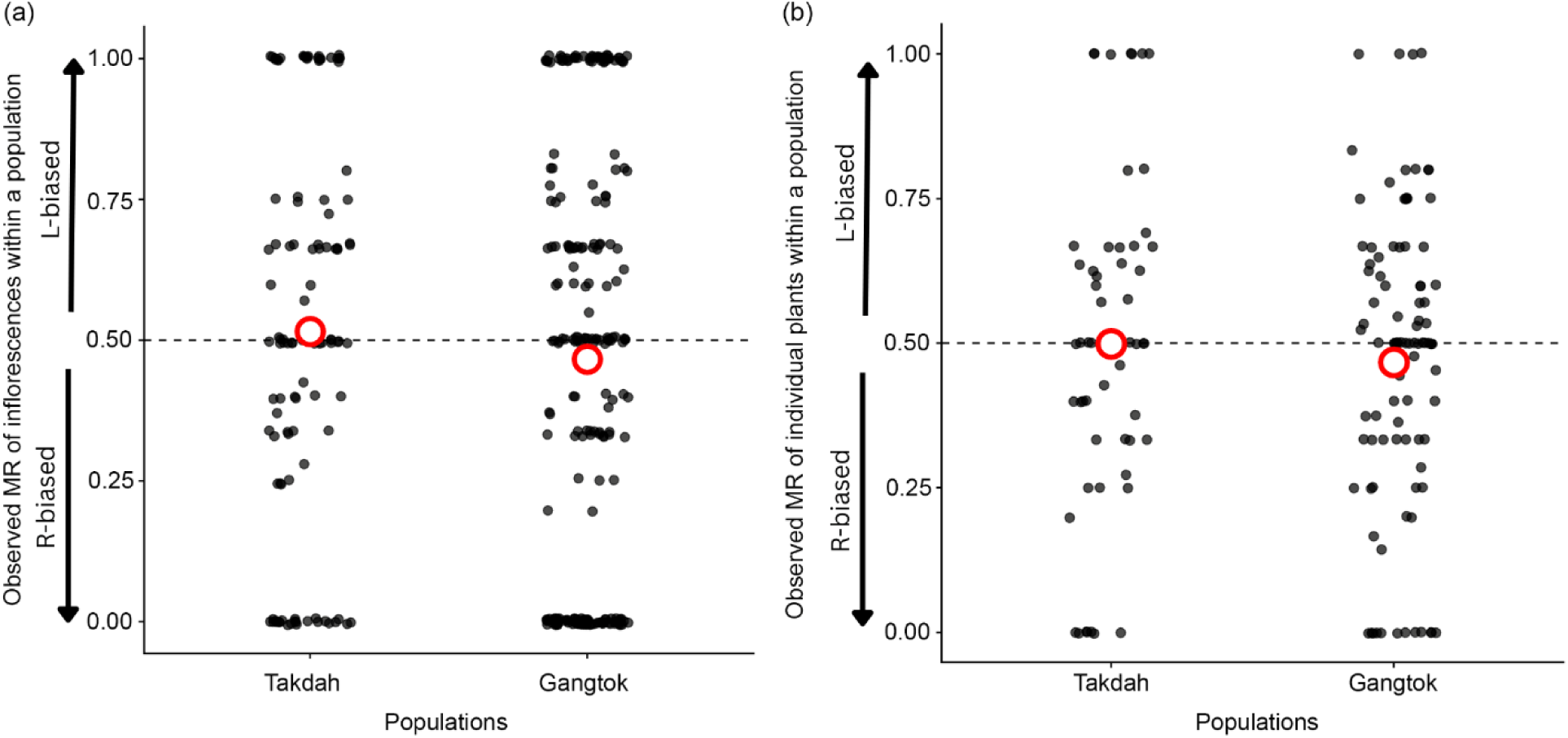
Observed morph ratio distributions in (**a**) inflorescences and (**b**) individual plants within two populations of *D. podocarpus*. Zero value on the y-axis represents R-bias, and one represents L-bias, while 0.5 represents an absolute isoplethic ratio. The red open circle represents the average of all the morph ratios distributed within the population. When the average value (red open circle) is at 0.5, it suggests that the population displays an equal number of L and R morphs (Gangtok population), while a slight shift towards the lower part of the isoplethic region suggests a slightly higher number of R morphs in the population (Takdah population). The data points here include single-flowered inflorescences and individuals.

### Estimation of morph ratios (MR) to quantify L bias and R bias

In ME, since both morphs can be present on the same inflorescence, an inflorescence can show L bias, R bias, or an isoplethic ratio. To estimate the relative morph abundance within an inflorescence, within an individual, and within a population, we sampled 30–33 individual plants at random, across three years in Takdah, in 2022 (T22), 2024 (T24), and 2025 (T25), and two years in Gangtok in 2024 (G24) and 2025 (G25). For each individual, we counted the number of open L and R morphs on all the inflorescences. Next, to quantify the relative abundance of the morphs within inflorescences and individual plants, we calculated the ‘morph ratio’ (MR) of inflorescences and individual plants as: MR = the total number of flowers of L morph ÷ the total number of flowers. Thus, MR represents the relative bias of inflorescences or plants with reference to only the L morphs and is a metric that retains morph identity (L bias or R bias). That is, when all flowers in an inflorescence or plant are L morphs, the MR value = 1, and can be identified as L bias. Later, we introduce another metric, morph bias (MB), which designates the overall morph bias of an inflorescence or plant, without retaining morph identity. To investigate yearly variation in the MR of plants, we fitted a Generalised Linear Mixed Model (GLMM) with a binomial distribution and logit link function using the lme4 package (Bates et al., 2015) in R (version 4.4.0; R Core Team, 2024). We used the number of L morphs within individual plants as the response variable with the total number of flowers as a binomial denominator, the year of sampling was the fixed effect, and the population was the random effect. The significance of the individual fixed effects was tested using Wald’s χ^2^ test (type III) using the car package (Fox & Weisberg, 2018), and multiple contrast analysis was carried out using the emmeans package (Lenth, 2025) with Sequential Bonferroni Correction. A chi-square goodness-of-fit test was used to evaluate whether the total number of L and R morph flowers within populations across years deviated from a 1:1 expectation.

To test whether the observed morph ratios within individual plants deviate from a stochastic expectation given the number of open flowers per plant, we used a maximum likelihood approach. Using this approach, we modelled the number of L morphs per plant given the total number of open flowers per individual. We compared two models: a binomial model assuming a fixed probability of occurrence of L morphs and a beta binomial model, where the underlying probability of morph occurrence varied between individual plants, thereby accounting for extra-binomial variation. The beta-binomial model was fitted using the VGAM package (Yee, 2015). Model fit was compared using the Akaike Information Criterion (AIC) values, with a lower value indicating better model support. All statistical analyses were carried out in R (version 4.4.0; R core team, 2024).

### Reproductive compatibility between morphs

To quantify the self and cross-compatibility between the two morphs, we carried out the following hand-pollination treatments: (i) inter-morph outcrossing (n = 40 flowers from 29 plants), (ii) intra-morph outcrossing (n = 41 flowers from 30 plants), (iii) inter-morph geitonogamy (n = 42 flowers from 25 plants), (iv) intra-morph geitonogamy (n = 40 flowers from 25 plants), and (v) unmanipulated, bagged flowers to check for autonomous self-pollination (n = 52 flowers from 34 plants). Flowers were bagged pre– and post-treatment to avoid pollinator visitation, emasculated before anther dehiscence, and hand-pollination was carried out within the first 18 hours post-anthesis. Each treatment was carried out on flowers of different plants, with no plant receiving more than one treatment. A maximum of two flowers per plant was used for any given treatment. For outcrossing, mixed pollen from multiple plants at least 3 m apart was used. Mature fruits, except the ones that were herbivored, were collected after three weeks for seed count analysis.

The seeds of *D. podocarpus* are tiny – *c.* 0.3 × 0.1 mm and are reported to be ‘numerous’. Therefore, seed counting for mature fruits was carried out using ImageJ software (Rueden *et al*., 2017). All seeds were carefully removed from the capsule and placed in a white weighing boat. High-resolution images of these weighing boats were taken using a digital camera (Nikon D3500), and the total number of seeds per capsule was quantified using the ‘analyse particle’ function in the ImageJ software following the method mentioned in Ochogavía (2022). This was carried out for a total of 84 fruits: (i) inter-morph outcrossing (n = 25 from 22 plants), (ii) intra-morph outcrossing (n = 21 from 18 plants), (iii) inter-morph geitonogamy (n = 23 from 17 plants), (iv) intra-morph geitonogamy (n = 15 from 12 plants).

We fitted separate GLMM models to check whether the fruiting success rate and seed count differed between (i) different hand pollination treatments, (ii) between L and R morphs, (iii) between outcrossing and selfing treatments, and (iv) between inter-morph and intra-morph treatments. Fruiting success (binary response: fruit set vs. no fruit set) was analysed using a binomial GLMM with logit link function, and seed count (seed count from all matured fruits as response variable) was analysed using a Poisson distribution with log link function. Seed counts between L and R morphs were compared using a negative binomial distribution with a log link function due to overdispersion and the poor fit of the Poisson distribution. Plant identity and population were treated as random effects in all models. The significance of the individual fixed effect and multiple contrast analysis was carried out using the method mentioned under the section ‘estimation of morph ratio, and to quantify L and R bias’.

Furthermore, to compare the pollen production between the two morphs, we counted the pollen grains from mature buds of both morphs. Paired mature buds (one L and one R morph) were collected from 30 individual plants (20 from Takdah and 10 from Gangtok). Undehisced anthers were stored in microcentrifuge tubes containing 1 ml of 70% ethanol. The anthers were macerated, and pollen grains were counted using a Neubauer chamber following the method mentioned in Dafni (1992) and Kearns & Inouye (1993). The difference in pollen count was compared using a negative binomial GLMM with log link function using glmmTMB (Brooks *et al*., 2017), with pollen count as response variable, morph identity as fixed effect and plant identity and population treated as random effects.

### Pollen placement on the pollinator

To examine if the pollen from the two morphs is differentially placed on the *Bombus* bees, we carried out controlled manipulative experiments using stained pollen grains and euthanised bees. The pollen grains of the L and R flowers were labelled using heavy-metal-free CuInSexS2-x/ZnS (core/shell) quantum dots with Zinc Oleate ligands dissolved in hexane (5 mg/mL), following the method mentioned in Minnaar and Anderson (2019). To distinguish the pollen grains of L and R morphs, we labelled them using 2 μL of yellow (580 nm) and red (650 nm) quantum dots, respectively. Experiments were carried out on 10 freshly opened flowers of each morph, and the individual euthanised *Bombus* bees (*n* = 10) were manually inserted into a labelled L morph flower, followed by an R morph using sterile forceps, mimicking their natural probing behaviour (Fig. 5a). To quantify the number of labelled pollen grains deposited on the pollinator, we categorised the body of the bees into four parts (Fig. 5a): (i) left proximal region (left side of head and thorax), (ii) left distal region (left side of the abdomen), (iii) right proximal region (right side of the head and thorax), and (iv) right distal region (right side of the abdomen). The number of fluorescent pollen grains was counted in these four regions using a 60X hand microscope with an inbuilt UV light source. We tested for differences in pollen placement on pollinators using a GLMM with a negative binomial distribution and log link function with the number of fluorescent labelled pollen grains as the response variable, the interaction between morph type and bee body part as a fixed effect, and bee identity as a random effect. The significance of the fixed effects and multiple contrast analysis was carried out as mentioned in the section ‘Estimation of morph ratios (MR) to quantify L bias and R bias’.

### Pollen movement between morphs using quantum dots

To confirm if pollen transfer to stigmas is highest between the two morphs (L to R and R to L) in natural conditions, we tracked the pollen using quantum dots. We selected 12 plots (eight from Takdah and four from Gangtok) of 3 × 2 m with a minimum distance of 1.5 km between plots. To mimic the natural ratio of morphs within a population in our experimental plots, we maintained the morph ratio at an isoplethic ratio (a 1:1 ratio of L and R morphs) by manually removing flowers when necessary. Since *D. podocarpus* has a restricted distribution in its native habitat and the available flowering density allowed only a single experimental setup, in the interest of the species and its conservation, we did not set up alternate experiments, such as with variable morph ratios. All the inflorescences within the plot were bagged a day before the quantum dot application to avoid pollen loss. To ensure recording of pollen movement in natural conditions, pollen grains of approximately one-fourth of the total number of L and R flowers were labelled between 06:00 and 07:00 h using yellow (for L-morph) and red (for R-morph) colored quantum dots. After 8 hours of natural pollination, all flowers within the plot were collected between 15:00 and 16:00 h and immediately screened for the number of fluorescent pollen grains deposited on the stigmas using a 60X hand microscope with an inbuilt UV light source. We tested if inter-morph pollen transfers were higher than intra-morph pollen transfer by fitting a negative binomial GLMM and a log link function, with the number of fluorescent labelled pollen grains on the stigma as a response variable, the interaction between morph type of the donor and recipient as a fixed effect, and plant and plot identity as random effects. The significance of the individual fixed effect was tested, and pairwise comparisons were carried out using the method mentioned in the section ‘Estimation of morph ratios (MR) to quantify L bias and R bias’.

### Quantifying pollinator constancy towards a floral morph

Pollinator preference for a morph and its subsequent foraging behaviour can influence pollen movement between morphs. To explore if individual pollinators show a preference for one morph while foraging across multiple plants, which may result in biased morph switches, we quantified the floral constancy of *Bombus* bees towards a specific morph in a single foraging bout. We tracked individual *Bombus* bees (*n* = 50) for up to 10–30 consecutive floral visits, and recorded their total intra-morph switches (indicating high pollinator constancy towards a morph) and inter-morph switches (indicating low pollinator constancy towards a morph) during a single foraging bout. We used a paired-sample t-test to assess whether the frequency of intra-morph switches significantly differed from that of inter-morph switches.

### Effect of morph bias on pollinator movement within the plant

Complementary to the above experiment, we next tested whether pollinator switches within a plant may be governed by the plant’s morph bias. For this, we first measured pollinator switches within a plant in 72 of the 153 experimental plants (40 in Takdah and 32 in Gangtok). Each plant was observed for two 25-minute periods from 07:00 to 15:00 h during the peak pollinator activity time, resulting in a total of 60 hours of observation (2 slots × 25 min × 72 plants). We recorded 76 *Bombus* bees and their within-plant inter-morph switches (R to L and L to R) and intra-morph switches (R to R and L to L) during one visitation bout. We next assigned a morph bias value for each experimental plant, where ‘morph bias’ (MB) was computed as: MB = 1-(number of flowers of the rare morph ÷ number of flowers of the abundant morph). Here, MB = 0 would represent individuals with equal proportions of both morphs, while MB =1 would indicate a complete skewness towards one morph. This metric identifies only if a plant deviates from an isoplethic morph ratio and does not identify the type of bias, that is, L bias or R bias. In ME, since both morphs are present on a plant, and all inter-morph switches (L to R and R to L) will result in self-pollen transfer (geitonogamy), the direction of the switch is not critical. To examine the effect of morph bias on the frequency of inter-morph switches, we fitted a GLMM with a binomial distribution and logit link function, with the number of inter-morph switches during each visitation bout as a response variable with total number of switches as a binomial denominator, MB as a fixed effect and plant identity and population as random variables.

Finally, we quantified the effect of MB on the pollination success of plants by counting the total number of fruits, normalised by the total number of flowers per plant, in a total of 109 (out of 153) naturally pollinated plants. To test the effect of morph bias on the fruit set, we used a binomial GLMM and logit link function, with the number of fruits per individual plant as a response variable, the total number of flowers as a binomial denominator, MB as a fixed effect and population as a random variable.

## Results

### Morph ratio (MR) within populations

We calculated the MR of a total of 338 inflorescences from 153 plants, from two populations (Table S1). The GLMM analysis revealed that the MR of plants across the years and for the two populations was not significantly different (χ² = 5.4, df = 4, *p* = 0.25; Fig. S2), and therefore, in all subsequent analyses, population identity has not been retained. Our analysis comparing the binomial and beta-binomial stochastic models showed nearly identical likelihoods (binomial: logLik = –221.59, AIC = 445.19; beta-binomial: logLik = –221.59, AIC = 447.19). The beta-binomial model did not improve the model fit (ΔAIC = 2), indicating no evidence of extra-binomial variation in morph ratios among individual plants.

Among the 338 inflorescences examined, 30.47% inflorescences had single flowers. Out of the remaining multi-flowered inflorescences (235 inflorescences), 23.83% exhibited an isoplethic morph ratio where L and R morphs are present in equal numbers, and 76.17% exhibited some deviation from isoplethy where one morph is greater in number than the other. Similarly, among the 153 individual plants examined, 5.88% plants had single flowers. Out of the remaining multi-flowered individual plants (144 individuals), 20.83% exhibited an isoplethic morph ratio, whereas 79.17% exhibited some deviation from isoplethy. However, within a population, the total number of L and R morph flowers across the years did not significantly deviate from an isoplethic ratio (T22: χ² = 3.47, *p* = 0.06; T24: χ² = 1.38, *p* = 0.24; T25: χ² = 0.04, *p* = 0.85; G24: χ² = 0, *p* = 1; G25: χ² = 0.59, *p* = 0.44; Table S1).

### Reproductive compatibilities of morphs

We carried out a total of 215 hand pollinations, of which 103 sired fruits. The per cent fruiting success of each pollination treatment was recorded as follows: (i) inter-morph outcrossing: 80%, (ii) intra-morph outcrossing: 61%, (iii) inter-morph geitonogamy: 66.7%, (iv) intra-morph geitonogamy: 45%, and (v) autonomous self-pollination with zero success rate (Fig. 3a). We observed a significant effect of the type of pollination treatment (excluding the autonomous self-pollination) on the fruiting success (χ² = 8.33, df = 3, *p* < 0.05), but the pairwise comparisons with sequential Bonferroni correction showed significant differences in fruit set only between inter-morph outcrossing and intra-morph geitonogamy (Table S2). The two morphs did not differ in their fruiting success (Estimate = –0.29, z = –0.8*, p* = 0.42). A marginally significant difference in fruit set was observed when all outcrossing treatments were compared with all selfing treatments, which included both intra and inter-morph crosses (Estimate = *-*0.75, z = –1.8*, p* = 0.07). However, the fruit set of all inter-morph pollination treatments was significantly higher than all intra-morph pollination treatments (Estimate = –0.99, z = –2.44*, p* = 0.01; Fig. 3a), irrespective of whether these crosses were carried out within a plant (selfing) or between plants (outcrossing).

**Fig. 3.**
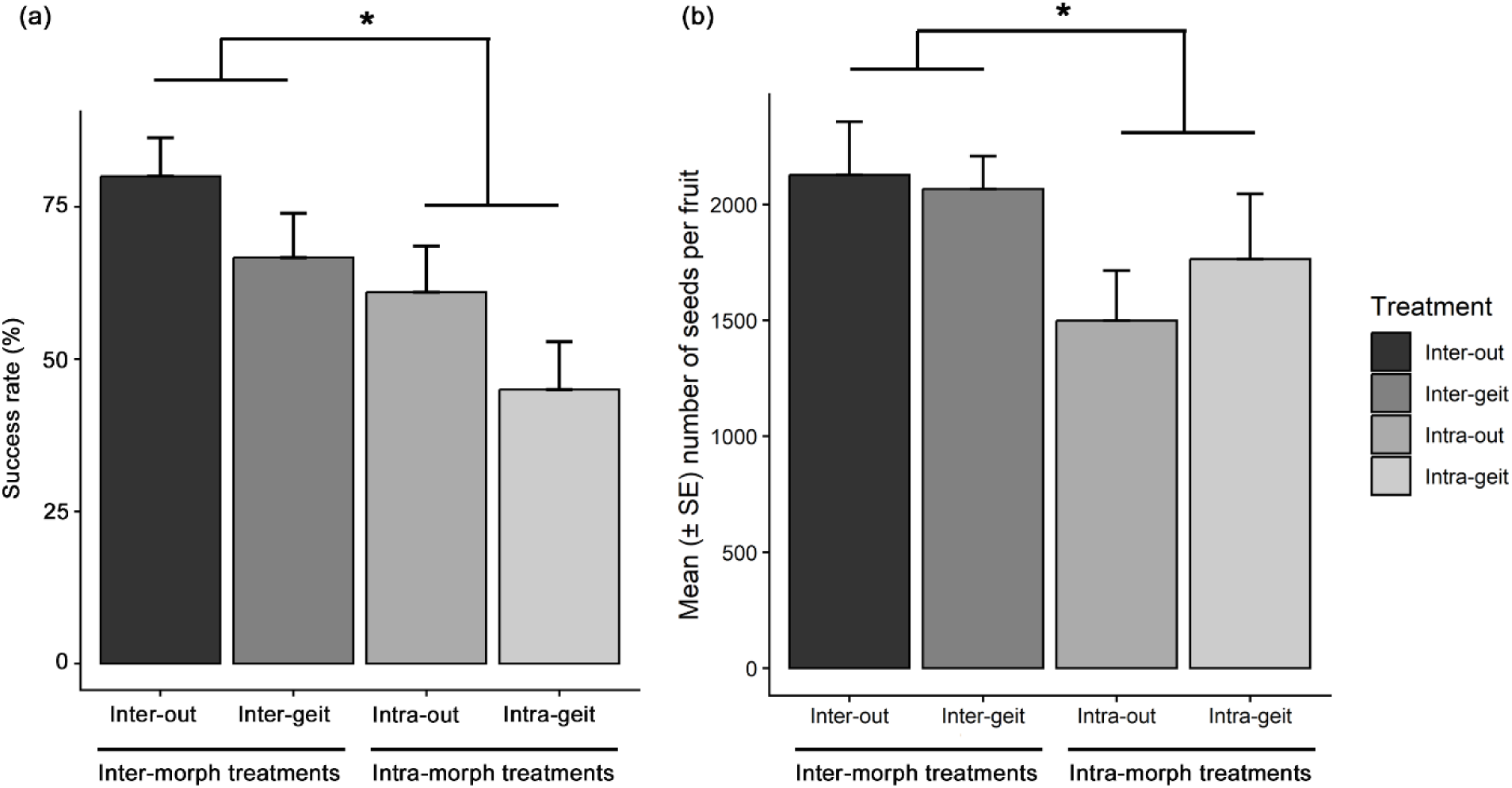
Results of fruiting success and seed count from four pollination treatments: inter-morph outcrossing (inter-out), inter-morph geitonogamy (inter-geit), intra-morph outcrossing (intra-out), and intra-morph geitonogamy (intra-geit). (**a**) Fruiting success rate showing a significant effect between inter-morph treatments and intra-morph treatments (inter-out: n = 40, inter-geit: n = 42, intra-out: n = 41, intra-geit: n = 40 flowers). Fruiting success was analysed using a binomial GLMM. Error bars represent standard errors calculated assuming a binomial distribution. (**b**) mean number of seeds per fruit (± SE) showing a significant effect between inter-morph treatments and intra-morph treatments (inter-out: n = 25, inter-geit: n = 23, intra-out: n = 21, intra-geit: n = 15 fruits). Seed count was analysed using a Poisson GLMM. * indicates a statistical significance at *p* < 0.05.

We did not observe a significant effect of the type of pollination treatment on seed count (χ² = 5.41, df = 3, *p* = 0.14; Fig. 3b), and it did not differ between the two morphs (Estimate = – 0.03, z = –0.19*, p* = 0.85). No significant difference was observed between all types of outcrossing and selfing treatments (Estimate = 0.1, z = 0.44*, p* = 0.66). However, consistent with the fruiting success, all inter-morph treatments produced significantly higher seeds compared to intra-morph treatments (Estimate = –0.5, z = –2.26*, p* < 0.05; Fig. 3b). Finally, pollen count was also noted to be different between the morphs, where flowers of the R morph show significantly higher pollen count when compared to the L morph (Estimate = 0.24, z = 3.81*, p* < 0.001; Fig. 4a).

**Fig. 4.**
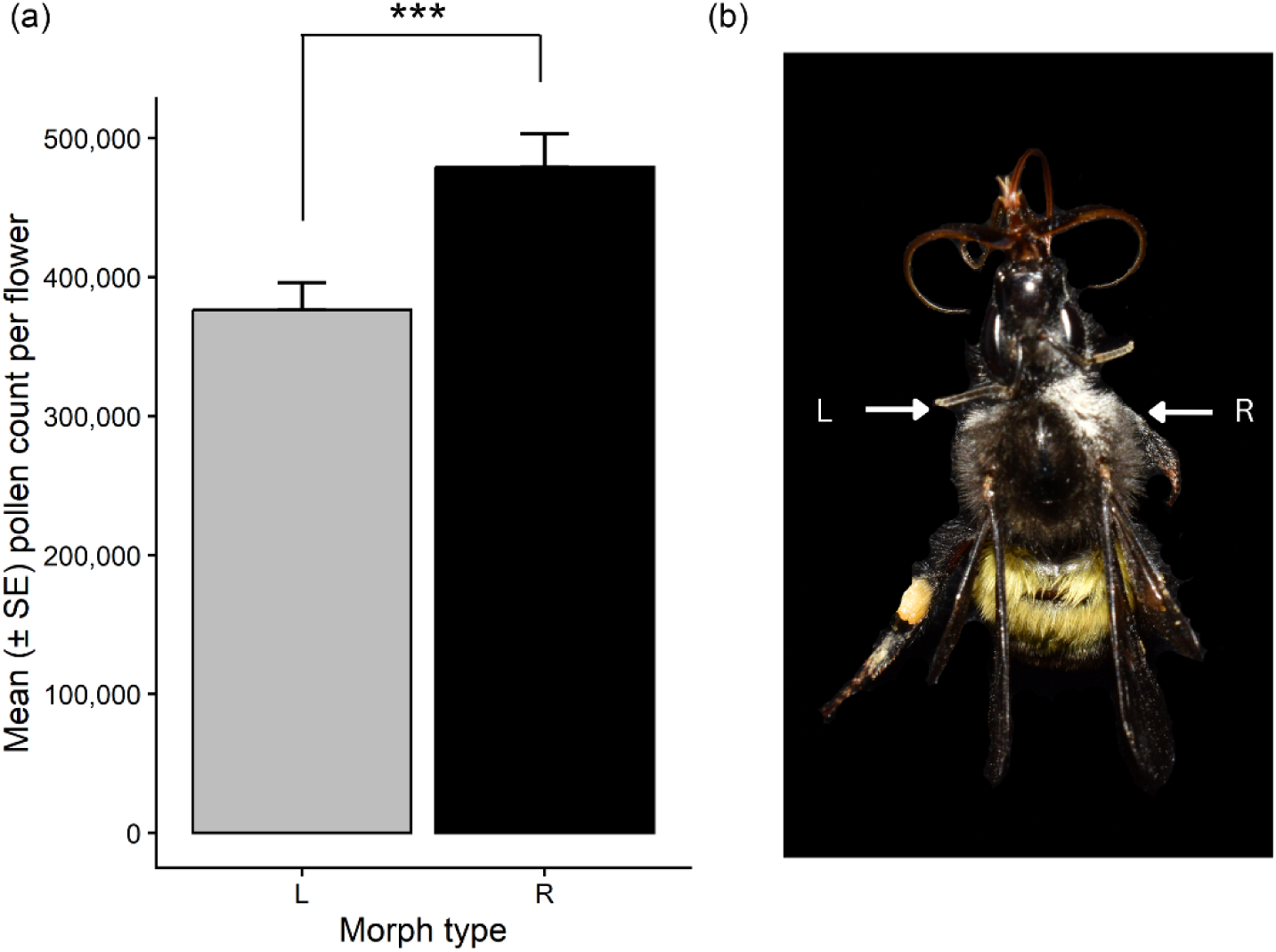
Comparative pollen count in the two morphs and on the pollinator. (**a**) Comparison of pollen count (mean ± SE) between L and R morph flowers from 30 individuals, analysed using a negative binomial GLMM. * indicates a statistical significance at *p* < 0.001. (**b**) Asymmetrical pollen deposition was noted on a *B. breviceps* after natural visits to approximately 30 flowers on different *D. podocarpus* plants. Arrows point to the larger pollen patch on the bee’s right side of the thorax, the contact area of the R morph (R), while the smaller pollen patch on the bee’s left side represents the contact area of the L morph (L).

### Pollen placement on the pollinator

All *B. breviceps* individuals (n=50) observed during their natural visitation to *D. podocarpus* probed the flowers using their proboscis by positioning their proximal part (head and thorax) at the rim of the flower. Since the anther and stigma are situated at the rim of the floral opening, the anthers of the L and R morphs were observed to come in contact with the left and right proximal regions of the Bombus bees, respectively (Fig. 4b). This observation was further corroborated by results from the pollen placement experiment using euthanized bees and pollen stained using quantum dots which show that the left and right proximal regions of the bees received pollen from the two morphs differentially (χ² = 75.65, df = 3, p < 0.001; Fig. 5b). The following pair-wise comparison shows that the left proximal region of the *Bombus* bee received a significantly higher number of L morph pollen grains than the R morph (Estimate = 2.6, z = 4.89*, p* < 0.001), and the right proximal region received more R morph pollen grains than the L morph (Estimate = –3.54, z = –6.64*, p* < 0.0001; Fig. 5b). We further found that the R morph deposited significantly higher labelled pollen grains on the pollinator compared to the L morph (χ² = 6.68, df = 1, *p* < 0.01).

### Pollen movement using quantum dots

All 12 plots in which pollen tracking experiments were carried out consisted of a total of 582 flowers, in which we found movement of labelled pollen grains in stigmas of 217 flowers (37.28%; Table S3). Inter-morph pollen transfer identified on the stigmas was significantly higher than the intra-morph pollen transfer (χ² = 126.3, df = 1, *p* < 0.0001; Fig. 6a). The difference in inter-morph pollen transfer was true when the L morph was the recipient flower (that is, R × L > L × L, donor × recipient; Estimate = –2.35, z = –10.86*, p* < 0.0001) as well as when the R morph was the recipient flower (that is, L × R > R × R; Estimate = 1.01, z = 4.99*, p* < 0.0001). However, irrespective of intra– or inter-morph comparisons, stigmas of the L morph received significantly higher labelled pollen than the R morph (χ² = 59.12, df = 1, *p* < 0.0001; Fig. 6). Additionally, donor morph also had a significant effect on pollen transfer (χ² = 118.03, df = 1, p < 0.0001), with pollen originating from R morph flowers contributing significantly more to overall pollen movement.

**Fig. 5.**
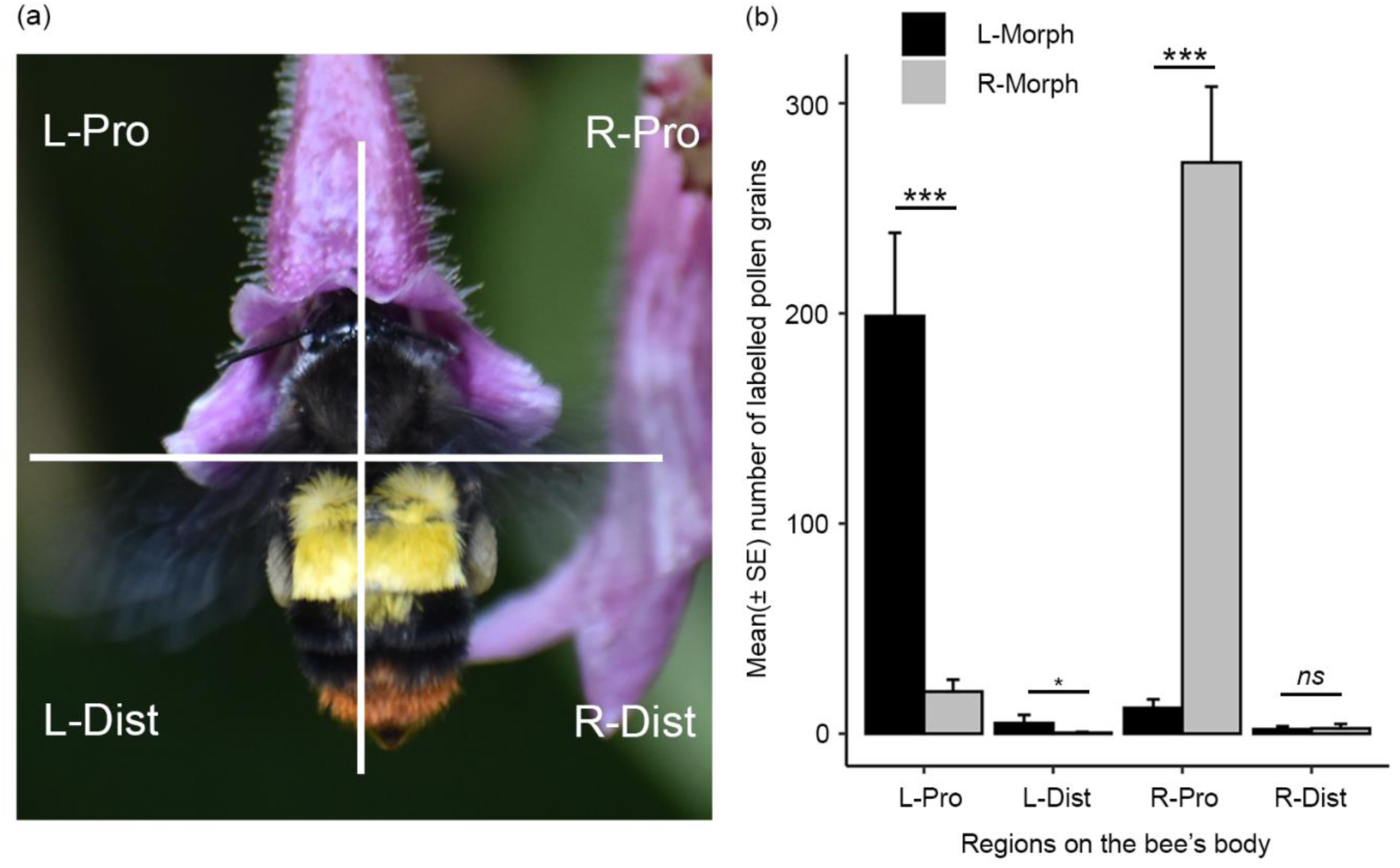
Pollen deposition across different parts of *B. breviceps*. (**a**) The natural probing position exhibited by *B breviceps* and the four regions in which pollen deposition was quantified: left proximal (L-Pro), left distal (L-Dist), right proximal (R-Pro), and right distal (R-Dist). (**b**) Mean (± SE) number of pollen grains labelled with quantum dots recovered from the euthanised *B. breviceps* after manual probing (n = 10 bees). Pollen counts were analysed using a negative binomial GLMM. Asterisks indicate statistical significance: ***, *p* ≤ 0·001; *, *p* ≤ 0·05; ns, non-significant.

**Fig. 6.**
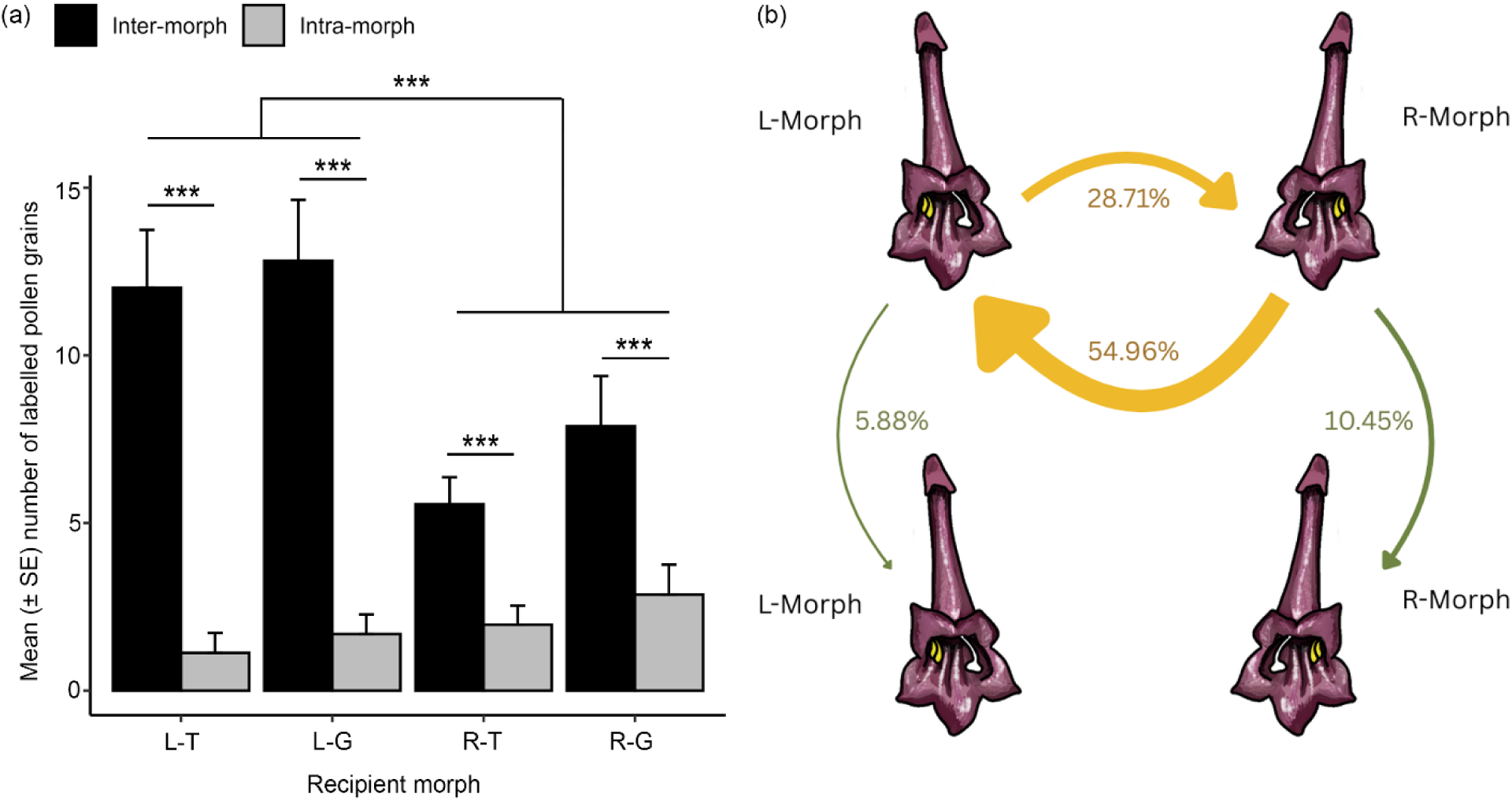
Pollen deposition on the stigmas of the two morphs in Takdah (T) and Gangtok (G) population. (**a**) Mean (± SE) number of inter-morph (black) and intra-morph (grey) quantum dot-labelled pollen grains deposited on the stigma of L and R morph recipients (n = 217 flowers). Pollen deposition on the stigma was analysed using a negative binomial GLMM. *** indicates statistical significance at *p* ≤ 0·001. (**b**) A schematic representation of the percentage of pollen movement recorded within morphs and between morphs of *D. podocarpus.* Illustrations by Abhishek Thakur.

### Pollinator constancy and pollinator movement within a plant

Paired t-test results did not show any significant difference between the number of intra-morph switches (pollinator constancy on a morph) and inter-morph switches (absence of pollinator constancy) by *Bombus* bees (*n* = 50) during their single foraging bout (*t* = 0.53, df = 49, *p* = 0.6). We noted a significant negative correlation between the frequency of inter-morph switches and MB (Estimate = –2.54, z = –3.69, *p* < 0.001; Fig. 7a). Specifically, pollinators exhibited higher inter-morph switches when the morphs were in equal proportion (isoplethic), and the frequency of these switches decreased as the morph bias approached one (complete bias, Fig. 7a). Finally, GLMM analysis also identified a significant positive association between the MB and the fruit set (Estimate = 0.35, z = 3.48, *p* < 0.001; Fig. 7b). That is, individual plants with a biased morph ratio exhibited higher fruit sets.

**Fig. 7.**
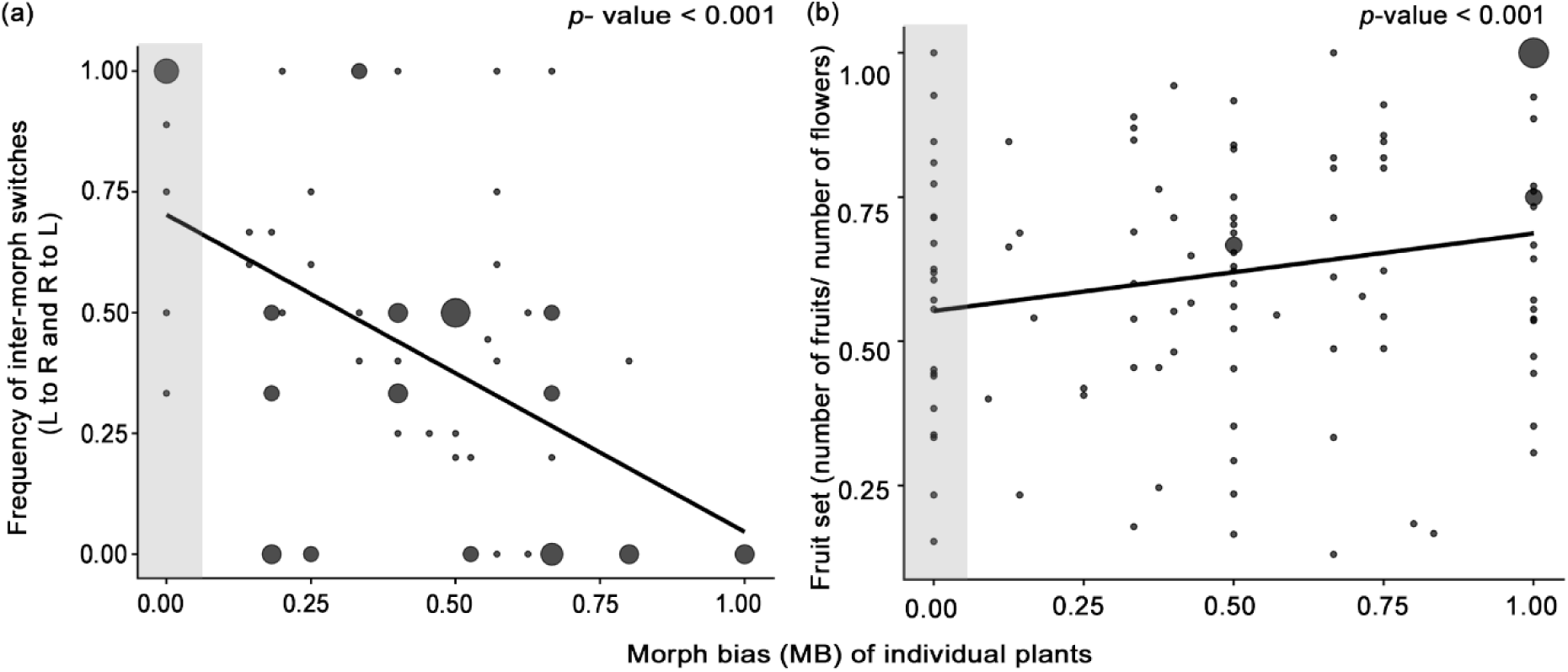
Relationship between morph bias (MB) and pollinator switches, and MB and fruiting success. (**a**) Correlation between the morph bias of individual plants and the frequency of inter-morph switches by the pollinator (n = 72 individuals), analysed using a binomial GLMM. (**b**) Correlation between the morph bias of individual plants and the natural fruit set (n= 109 individuals), analysed using binomial GLMM. The values range from 0 (isoplethic), which represents an equal number of both morphs, to 1, which represents complete skewness towards either one of the morphs. The grey shaded area along the x-axis represents plants with isoplethic morph ratio. Statistical significance is indicated in the top right corner. The size of each circle represents the number of observations for each combination of morph bias and the corresponding response variable. In (a), circles represent the frequency of inter-morph switches for the respective morph bias, with the smallest circle indicating a single observation and the largest indicating seven observations. In (b), circles represent fruiting success (number of fruits/total number of flowers per individual) for the respective morph bias values, with the smallest circle indicating a single observation and the largest indicating six observations.

## Discussion

The relative abundance of morphs within an individual plant and its effect on opportunities for selfing (geitonogamy) remain one of the least explored aspects of monomorphic enantiostylous plants. The key findings from our study are that the majority of inflorescences and individuals of the monomorphic enantiostylous *D. podocarpus* exhibited a biased morph ratio where one morph was dominant over the other, and this did not deviate from a binomial stochastic expectation given the total number of flowers per individual. However, the overall number of morphs within the population met an isoplethic ratio. We found that the frequency of inter-morph switches by the pollinator reduced with an increase in morph bias. Further, we report that inter-morph pollen transfers are higher than intra-morph pollen transfers, and hand-pollination treatments between morphs yielded a higher fruit set and seed set than hand-pollination treatments within morphs. Therefore, we propose that the predominant disassortative pollen movement and reduced inter-morph switches by the pollinators within individuals with higher morph bias may play a critical role in reducing self-pollination (geitonogamy) and increasing cross-pollination.

### Individuals with biased morph ratios show reduced geitonogamous events

The observed morph ratio within inflorescences and individual plants did not deviate from a binomial stochastic expectation, suggesting that the relative abundance of morphs in *D. podocarpus* may not represent a deterministic process at the individual level. We propose that the effect of a simple stochastic process in a system with small number of open flowers within individual plants (mean ± SE number of open flowers per individual plant is 6.28 ± 0.41) and inflorescences (mean ± SE number of open flowers per inflorescence is 2.84 ± 0.11) has resulted in higher number of plants within a population deviating from an isoplethic ratio, that is displaying morph bias, where one morph is present in greater number than the other (Table S1). Similar morph bias has also been reported from other ME species (Dulberger & Ornduff, 1980; Tang & Huang, 2005; De Almeida *et al*., 2018; Robertson *et al*., 2025). We show that the frequent morph bias observed within inflorescences and individuals (Table S1) in the two populations of *D. podocarpus* across multiple years has functional consequences. For example, in ME species, a morph bias within inflorescences and within individual plants can reduce the frequency of geitonogamous selfing events. This is because when there are multiple flowers on an individual plant, interactions specifically between flowers within an inflorescence can affect their reproductive outcome since pollinators tend to visit flowers within the same inflorescence, causing geitonogamy (Harder & Barrett, 1995; Harder *et al*., 2000; Harder *et al*., 2004). Therefore, any common inflorescence trait, such as the stochastic skew in individuals of the two *D. podocarpus* populations, is likely to influence pollination success and mating outcome (Harder *et al*., 2000).

Enantiostyly in *D. podocarpus* is unique because it differs from expected floral traits in enantiostylous flowers, such as the presence of heteranthery, the lack of nectary, and an open floral design (outward facing; Jesson & Barrett, 2003). Instead, *D. podocarpus* lacks heteranthery, has nectaries, and exhibits tubular flowers where the stigma and anther are exerted at the rim of the floral tube (Fig. 1). As expected from the floral morphology, and consistent with other studies, we found that inter-morph pollen transfers (L to R and R to L) were higher than intra-morph (L to L and R to R) pollen transfers, confirming disassortative pollen movement in *D. podocarpus* (Jesson & Barrett, 2005; De Almeida *et al*., 2013; Minnaar & Anderson, 2021; Fig. 6). In the quantum dot experiments to record pollen movement among and between morphs, overall we noted a smaller fraction of pollen deposited on the stigmas (Fig. 6) than what we observed on the pollinator in our manipulative experiment (Fig. 5). We propose that our observations have captured a realistic quantity of pollen deposition where the reduced pollen load may represent pollen loss due to pollinator grooming, also noted in other studies (Thompson, 1986), or pollen loss due to the dislodgment and loss of pollen from the stigma due to multiple pollinator visits (Stavert *et al*., 2020). Since we recorded pollen movement at the end of an 8-hour open-visitation period, we expect multiple pollinator visits to have occurred. Thus, a large pollen load that can be expected from a single virgin visit by a pollinator is absent in our quantum dot experiment.

Studies on heterostylous species have shown that, apart from the arrangement of sex organs within flowers, the floral display of individual plants and the morph composition of neighbouring plants affect the degree of disassortative pollination and quality of pollen deposited on the stigma (Waites & Ågren, 2004; Brys & Jacquemyn, 2010). While these factors are also true for ME species, the morph ratio within individual plants adds an additional layer of complexity. Results from our study provide empirical evidence of the theoretical prediction that the extent of geitonogamy that may occur in ME species depends on a) the relative abundance of L and R morph flowers within an individual plant, and b) the visitation sequence by the pollinator (Jesson & Barrett, 2005; Saltini *et al*., 2025). Using inter-morph switches by pollinator within individual plants as a proxy, we found that a biased morph ratio within individual plants could limit the occurrence of geitonogamous events, and thus may influence both the male and female fitnesses by limiting the effect of both pollen and ovule discounting. The potential outcome of reduced selfing was also reflected in the fruit set, where the fruit set increased with an increase in the morph bias.

We expected that the predominant pollen movement between flowers of different morphs might mandate an equal proportion of both morphs in a population. Our results validate this because, despite the biased morph ratio in the inflorescence and in the individual plants, the total number of L and R morph flowers within the two populations across multiple years did not significantly vary from the isoplethic ratio. This result is also consistent with the majority of previous studies on ME species (Fenster, 1995; Ren *et al*., 2013; Morais *et al*., 2020; Contreras-Varela *et al*., 2023). This suggests that, while individuals exhibit a biased morph ratio, the isoplethic ratio at a population level is likely driven by negative frequency-dependent selection since any deviation from this ratio can lead to lower pollination success (Jesson & Barrett, 2002).

### Flowers of the same morph show reduced compatibility

Studies examining reproductive compatibilities among morphs have reported that stylar polymorphisms, like distyly and tristyly, are linked to heteromorphic incompatibility associated with variations in pollen size and number, size of the stigmatic papillae, and stigma shape between morphs, potentially leading to intra-morph pollen recognition at the stigma, style, or ovary, thus reducing fruiting success and seed count (Richards, 1977; Barrett & Cruzan, 1994; Massinga *et al*., 2005; Costa *et al*., 2017). We noted moderate intra-morph incompatibility in the form of lower fruit set and lower seed count per fruit in *D. podocarpus,* irrespective of whether the intra-morph treatment was within or between plants. These observations strongly suggest that ‘morph type’ is being recognised regardless of whether the pollen is from within the individual or a different individual. Intra-morph incompatibility has been previously reported in the dimorphic enantiostylous species *Wachendorfia paniculata* (Ornduff & Dulberger, 1978), although a subsequent study failed to corroborate this (Jesson & Barrett, 2002). We propose that the moderate incompatibility between flowers of the same morphs in *D. podocarpus* can act as an additional mechanism to ensure disassortative pollen movement (Costa *et al*., 2017), where the fruiting success of pollen transfer between flowers of the same morph is significantly reduced.

Finally, in our experiments to track pollen movement, despite an equal number of L and R morphs in our experimental plots, the stigmas of L morphs received significantly higher numbers of labelled inter-morph pollen compared to stigmas of R morphs (Fig. 6). Similarly, R morphs deposited significantly higher pollen on the pollinator compared to the L morphs (Fig. 4b; Fig. 5b). In concordance with this, the comparison of pollen count between the two morphs show that the R morph indeed produces significantly higher pollen compared to the L morph in *D. podocarpus* (Fig. 4a). This difference was observed only when the morph was a pollen donor, and the fruiting success of the two morphs as a female parent was not different in our hand pollination experiments. These results suggest that the two morphs may have dissimilar sexual roles due to their differential contribution to the population’s overall pollen pool. The R morph in

*D. podocarpus* appears to display higher male function through higher pollen donation, whereas the L morph may display higher female function by receiving a higher number of pollen grains. These patterns loosely share similarities with a few andromonoecious systems, where male flowers are reported to produce more pollen compared to that of hermaphrodite flowers (Narbona *et al*., 2005; Dai & Galloway, 2012). While male flowers of andromonoecious plants are solely males, in the monomorphic enantiostylous *D. podocarpus* R morphs may emphasise more on male function by producing and exporting more pollen without being exclusively male.

## Conclusion

Monomorphic enantiostylous plants represent one of the unique innovations among labile reproductive strategies found in flowering plants, because, unlike other stylar polymorphisms, both morphs are found on the same plant. Here, we show that in the monomorphic enantiostylous *D. podocarpus*, the morph ratio is maintained at an isoplethic ratio in the population, even when the morph ratio within inflorescences and individuals remains highly variable. We also show that the stochastic skew in morph ratios within inflorescences and individuals, together with disassortative pollen movement and intra-morph incompatibility, appears to function as a multi-layered mechanism in reducing selfing opportunities (geitonogamy). Finally, we report intra-morph incompatibility along with the discovery of morph-specific differences in pollen count in the ME *D. podocarpus*, which is an intriguing result since morphs on the same individual share the same genetic background and yet displayed variable pollen counts. While this points to possible differences in the sexual roles of the two morphs on the same plant within an ME species, this is a topic that remains underexplored in plant reproductive biology. We believe that results from our ecological and behavioural studies will help future studies that may combine genetic, developmental and physiological tools to explore not only the differences between the morphs in the ME *D. podocarpus*, but also to understand their maintenance in nature. In conclusion, we show that a monomorphic enantiostylous plant enhances its opportunities for cross-pollination while also retaining reproductive assurance through the production of both morphs within a plant.

## Acknowledgements

We express our gratitude to the forest departments of Sikkim and West Bengal for the necessary permits. We would also like to thank the local community, especially Mr Provesh Gurung and his family, for their hospitality and support. We would like to thank Sanika Goray, Manila Chingtham, Rohan Dandavate, and Jyotil Dave for their assistance during the fieldwork, Kirti and Arti for helping with the seed count and Nevil S for assistance during the fieldwork and pollen count. We acknowledge Dr Paul Williams, Natural History Museum, London, and Dr A Rameshkumar, Zoological Survey of India, for helping us identify the *Bombus* bee specimens. We thank Dr N. S. Prasanna for giving us the details of the study sites. We thank Abhishek Thakur for the illustrations. We would like to acknowledge the Science and Engineering Research Board (SERB ECR/2017/001073; POWER Grant SPG/2021/000793) and DBT-NER (BT/PR24525/NER/95/754/2017) and Ministry of Human Resource Development (Ministry of Education), for the research fund awarded to VG, IISER Bhopal for infrastructural support, the University Grant Commission (UGC; 191620076099) for a fellowship to RSB and the American Society of Plant Taxonomists (ASPT) Graduate Student Research Grant, International Association for Plant Taxonomist (IAPT) Graduate Student Research Grant and Society for Tropical Ecology Student Research Grant awarded to RSB for fieldwork.

## Author contributions

RSB and VG conceived and designed the study. RSB carried out the field experiments and collected the data. RSB carried out the statistical analyses with inputs from VG. RSB drafted the manuscript, and VG provided conceptual advice and edited the manuscript. Both authors gave final approval for publication.

## Data availability

The data and R scripts used for this study will be made available in a public repository once the study is published.

