## Supporting information for "Morph bias in inflorescences and individual plants reduces opportunities for geitonogamy in a monomorphic enantiostylous species"

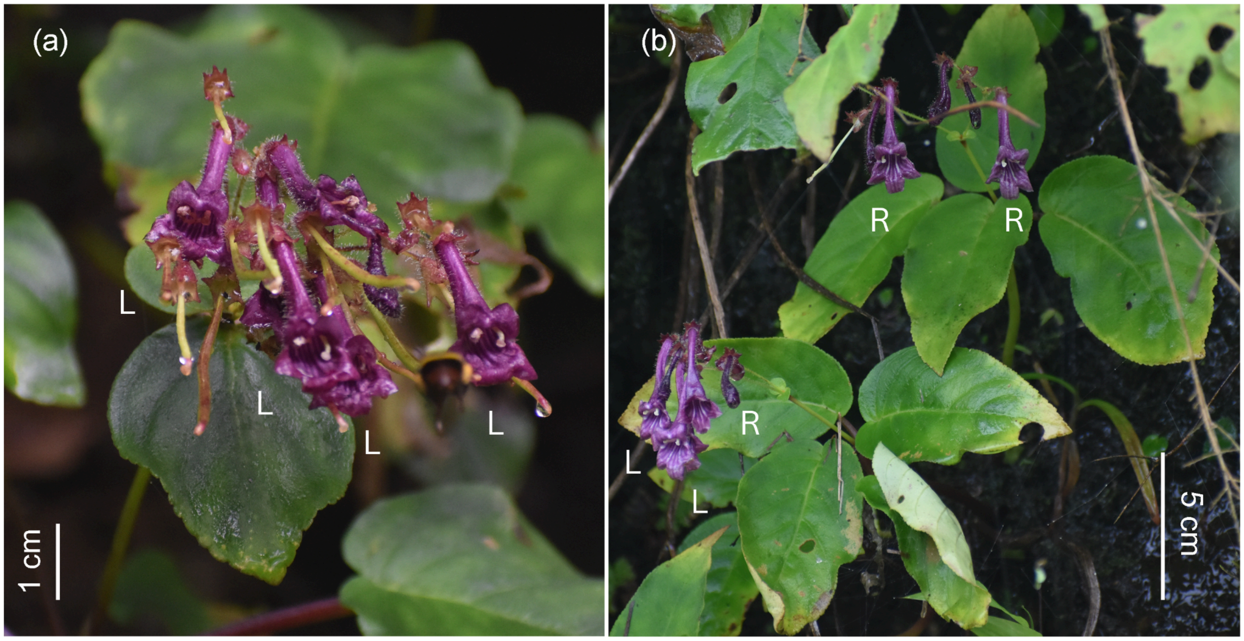


**Fig. S1** Image of *D. podocarpus* inflorescence and an individual plant with two inflorescences. (a) Arrangement of flowers of different morphs within an inflorescence, and (b) arrangement of inflorescences within an individual plant. Arrangement of flowers within an inflorescence and inflorescences within an individual plant do not show a predictable pattern. The two morphs are labelled as L and R, representing L-morph and R-morph flowers, respectively.

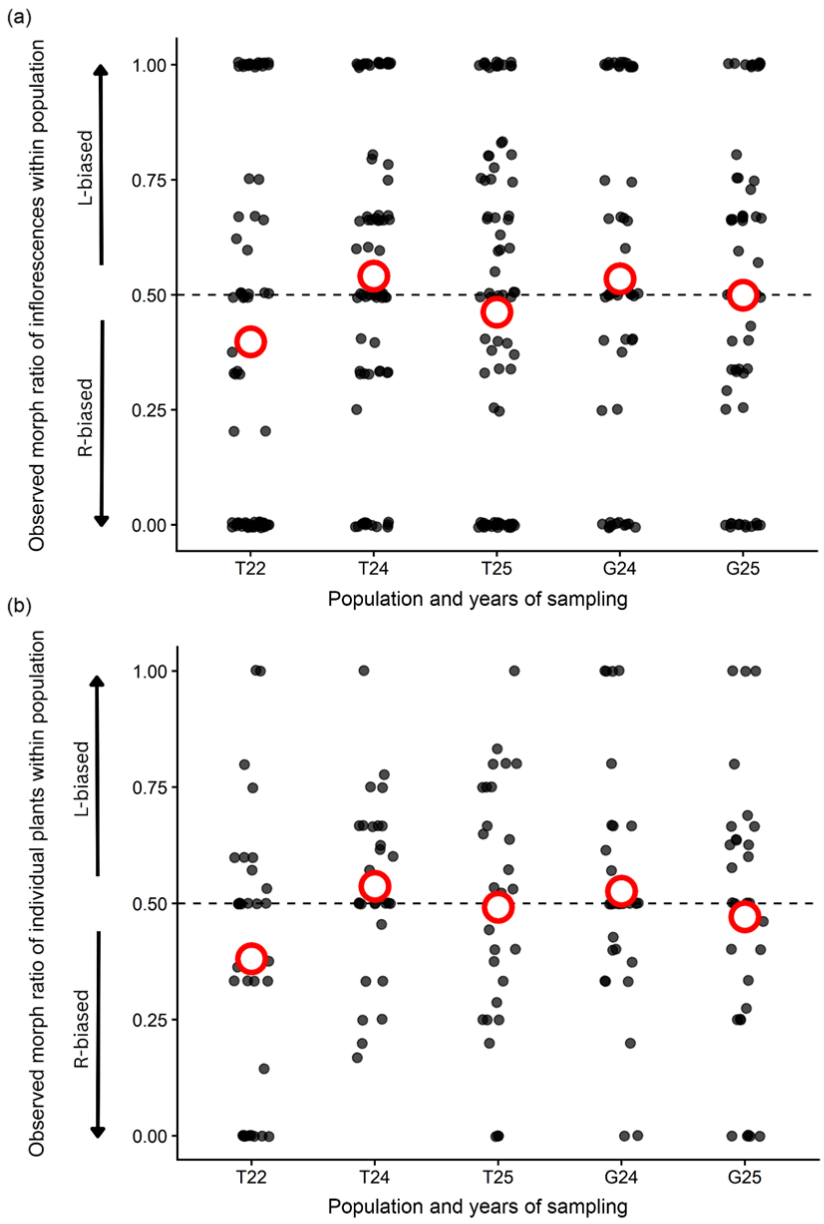


**Fig. S2** Observed morph ratio distributions in (a) inflorescences and (b) individual plants within two populations across multiple years of sampling. The x-axis represents the years of sampling in the Takdah population in 2022 (T22), 2024 (T24), and 2025 (T25), as well as the Gangtok population in 2024 (G24) and 2025 (G25). Zero value on the y-axis represents R-bias, and one represents L-bias, while 0.5 represents an absolute isoplethic ratio. Inflorescences and individual plants exhibiting an isoplethic morph ratio are represented along the dotted line. The red open circle represents the average of all the morph ratios distributed within the population. The data points here include single-flowered inflorescences and individuals.

| Population identity and sampling year | Total number of flowers | Number of L flowers | Number of R flowers | Number of plants with single flowers | Number of plants with isoplethic ratio | Number of plants with morph bias |
|---|---|---|---|---|---|---|
| T22 | 166 | 71 | 95 | 6 | 7 | 20 |
| T24 | 209 | 113 | 96 | 1 | 8 | 21 |
| T25 | 247 | 125 | 122 | 0 | 1 | 29 |
| G24 | 134 | 67 | 67 | 1 | 10 | 19 |
| G25 | 205 | 108 | 97 | 1 | 4 | 25 |

**Table S1** Total number of L and R morph flowers within a population and number of individuals with single flowers, isoplethic ratio, and morph bias (one morph greater in number than the other) during the time of census. The table includes observations from the Takdah population in 2022 (T22), 2024 (T24), and 2025 (T25), and the Gangtok population in 2024 (G24) and 2025 (G25).

| **Group 1** | **Group 2** | **Estimate** | ***p*-value** |
|---|---|---|---|
| Inter-geit | Inter-out | -0.8 | 0.5 |
| Inter-geit | Intra-geit | 1.01 | 0.34 |
| **Inter-out** | **Intra-geit** | **1.82** | **0.03*** |
| Inter-geit | Intra-out | 0.26 | 0.63 |
| Inter-out | Intra-out | 1.06 | 0.34 |
| Intra-geit | Intra-out | -0.75 | 0.5 |

**Table S2** Results of the pairwise comparison between four hand-pollination treatments: inter-morph outcrossing (inter-out), inter-morph geitonogamy (inter-geit), intra-morph outcrossing (intra-out), and intra-morph geitonogamy (intra-geit).

| Population | Plot ID | Total number of: | | Pollen transfer was observed in: | |
|---|---|---|---|---|---|
| | | **L flowers** | **R flowers** | **L flowers** | **R flowers** |
| Takdah | 1 | 11 | 11 | 10 | 6 |
| | 2 | 12 | 12 | 4 | 2 |
| | 3 | 24 | 24 | 12 | 16 |
| | 4 | 32 | 32 | 10 | 13 |
| | 5 | 36 | 36 | 4 | 4 |
| | 6 | 25 | 25 | 5 | 3 |
| | 7 | 14 | 14 | 9 | 11 |
| | 8 | 17 | 17 | 10 | 9 |
| Gangtok | 9 | 38 | 38 | 15 | 14 |
| | 10 | 24 | 24 | 11 | 7 |
| | 11 | 14 | 14 | 1 | 4 |
| | 12 | 32 | 32 | 18 | 19 |
| Total | 12 | 291 | 291 | 109 | 108 |

**Table S3** Experimental flowers used across a total of 12 experimental plots in two populations of *D. podocarpus*. The table lists details of the total number of flowers of both morphs within the experimental plot during quantum dot application, and the number of flowers in which quantum dot-labelled pollen grains were recorded after allowing natural visits within the experimental plots.
